# Graph Attention Neural Networks Reveal TnsC Filament Assembly in a CRISPR-Associated Transposon

**DOI:** 10.1101/2025.06.17.659969

**Authors:** Chinmai Pindi, Mohd Ahsan, Souvik Sinha, Giulia Palermo

## Abstract

CRISPR-associated transposons (CAST) enable programmable, RNA-guided DNA integration, marking a transformative advancement in genome engineering. A central player in the type V-K CAST system is the AAA+ ATPase TnsC, which assembles into helical filaments on double-stranded DNA (dsDNA) to orchestrate target site recognition and transposition. Despite its essential role, the molecular mechanisms underlying TnsC filament nucleation and elongation remain poorly understood. Here, multiple-microsecond and free energy simulations are combined with deep learning-based Graph Attention Network (GAT) models to elucidate the mechanistic principles of TnsC filament formation and growth. Our findings reveal that ATP binding promotes TnsC nucleation by inducing DNA remodelling and stabilizing key protein-DNA interactions, particularly through conserved residues in the initiator-specific motif (ISM). Furthermore, GNN-based attention analyses identify a directional bias in filament elongation in the 5′→3′ direction and uncover a dynamic compensation mechanism between incoming and bound monomers that facilitate directional growth along dsDNA. By leveraging deep learning–based graph representations, our GAT model provides interpretable mechanistic insights from complex molecular simulations and is readily adaptable to a wide range of biological systems. Altogether, these findings establish a mechanistic framework for TnsC filament dynamics and directional elongation, advancing the rational design of CAST systems with enhanced precision and efficiency.

---

CRISPR-associated transposons (CAST) are genetic elements that integrate CRISPR systems with transposase enzymes, enabling precise, RNA-guided DNA insertion at specific genomic sites^1,2^. Their discovery marks a transformative advance in life sciences, unlocking the potential for large-scale, targeted DNA integrations^3–8^. Elucidating the biophysical principles underlying CAST-mediated target recognition and DNA integration is both fascinating and critical to harnessing their full potential in genome engineering applications.

The type V-K CAST system employs an RNA-guided Cas12k protein, the transposition regulator TnsC, and target site recognition factors to direct the transposase to specific DNA integration sites (**Fig. 1a**)^9–16^. TnsC, a putative AAA+ (ATPases Associated with diverse cellular Activities) protein, acts as a critical molecular bridge between target site recognition and transposase recruitment. Recent structural and biochemical studies of the *S. hofmann i*CAST (ShCAST) have shown that, in the presence of ATP, TnsC polymerizes into a helical filament that wraps around double-stranded DNA (dsDNA) (**Fig. 1a-b**)^9,10^. This polymerization is essential for RNA-guided transposition, as it enables target site selection, connects the transposase with DNA, and facilitates the recruitment of additional transposition factors^15,17^. Emerging evidence also suggests that TnsC filament formation is also critical for RNA-independent transposition, driving untargeted DNA integration at random genomic loci^18^.

**Fig. 1.**
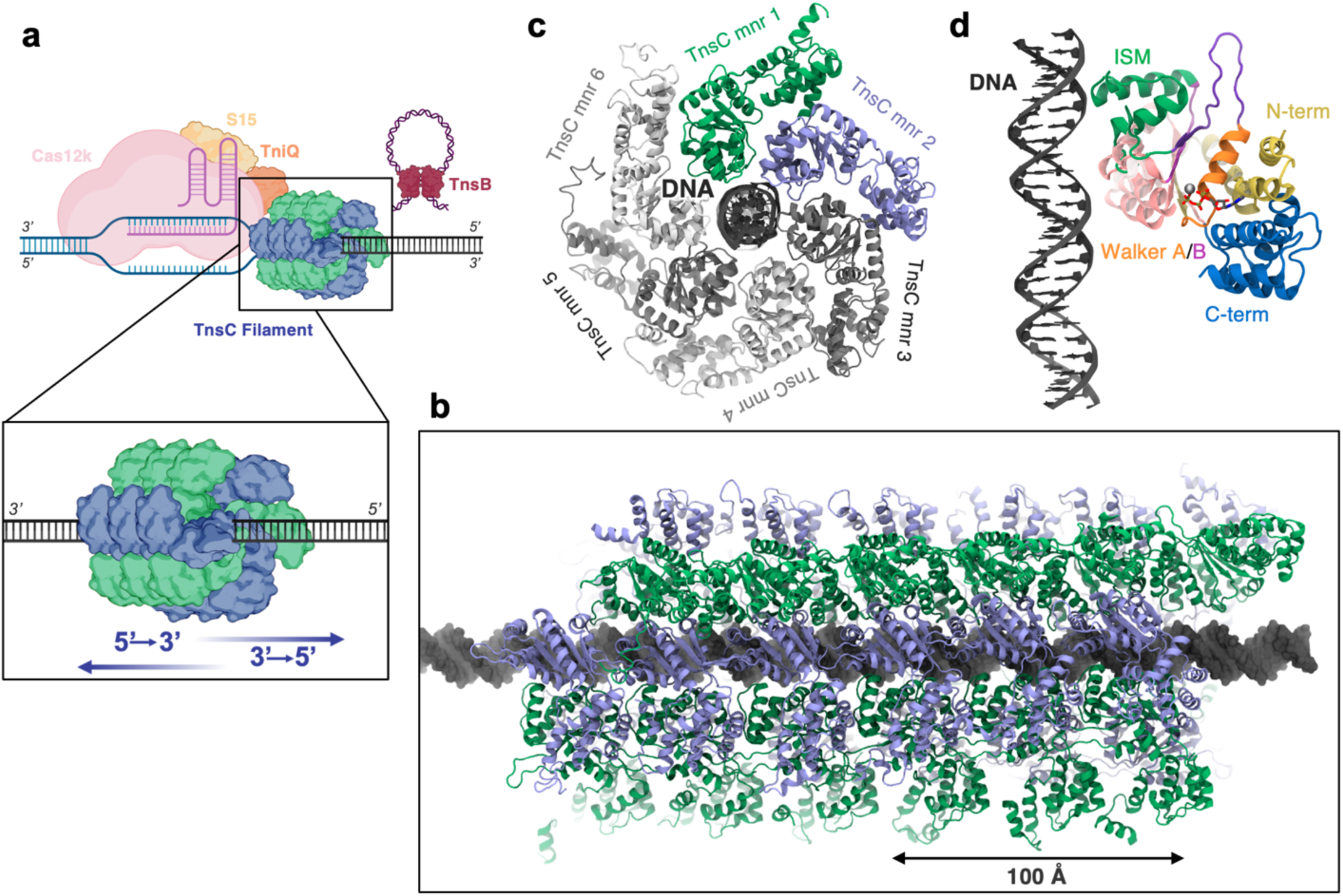
Overview of the type V-K *S. hofmanni* CRISPR-Associated Transposon (CAST) system. **a.** Schematic of the type V-K CAST transposition machinery, including the RNA-guided Cas12k protein and associated factors, the TnsC transposition protein polymerizing along double-stranded DNA (dsDNA), and the transposase enzyme. A magnified inset illustrates the two plausible directions of TnsC filament growth (i.e., 5′→3′ or 3′→5′). **b.** Structure of ~23 nm-long TnsC filament assembled on dsDNA, comprising six TnsC hexamers, reconstructed from the cryo-EM structure of the TnsC octamer (PDB 7M99)^9^. **c.** Each hexamer (top-view) includes six TnsC monomers that wrap around dsDNA. **d.** Close-up view of a single TnsC monomer bound to dsDNA, highlighting the bound ATP molecule (shown as sticks) and the coordinating Mg²⁺ ion (shown as a sphere). Individual subdomains of the TnsC monomer are color-coded (C-terminus: blue; N-terminus: yellow; Walker A: orange; Walker B: magenta; ISM: green; and two linker regions, *viz.*, linker 1 between Walker A and ISM: violet; linker 2 between Walker B and the C-terminus: pink).

In this context, TnsC’s ability to form filaments on dsDNA is emerging as a key driver of untargeted DNA insertions, enabling integration at random genomic sites. Despite its central role in transposition, the molecular mechanisms underlying TnsC nucleation and filament elongation remain poorly understood. Key unresolved questions include the molecular triggers that initiate TnsC polymerization in the presence of ATP, and the directionality of filament growth – whether it proceeds in the 5′→3′ direction toward the recognition complex or in the opposite 3′→5′ direction (**Fig. 1a**, close-up view). While structural studies have captured filaments in both directions ^14–16,18^, the biophysical factors that govern the directionality and dynamics of TnsC filament assembly remain elusive. Addressing these questions is essential for elucidating the molecular basis of TnsC-mediated integration and for advancing CAST systems as tools for precision genome engineering.

Structural studies have shown that ATP-bound TnsC polymerizes into spiralling helical filaments around dsDNA, with lengths ranging from ~22 to ~70 nm^9,10^. Electron microscopy and high-resolution structural data reveal a filament architecture with a pitch of ~40.7 Å and a rise of ~6.8 Å, comprising six TnsC monomers per turn (hereafter referred to as a TnsC hexamer, **Fig. 1c**). Each TnsC monomer features a canonical bipartite AAA+ ATPase domain comprising an α/β domain – with a central five-stranded β-sheet flanked by α-helices – and a C-terminal α-helical domain (**Fig. 1d**). The conserved Walker A and Walker B motifs form the ATP-binding pocket, flanked by the Initiator-Specific Motif (ISM), as well as the N- and C-termini and two linker regions (Linker 1 between Walker A and ISM; Linker 2 between Walker B and the C-terminus). This domain organization enables TnsC to adopt the spiral, corkscrew-like conformation characteristic of initiator-clade AAA+ proteins as they assemble around dsDNA^19^. Notably, unlike other non-CAST transposon-associated AAA+ proteins – such as MuB ^20^, which form helical oligomers in the presence of ATP alone – TnsC requires both ATP and DNA for filament formation DNA^9,10^. Understanding the mechanism of TnsC polymerization and filament assembly is thus critical not only for elucidating its role in CAST-mediated DNA transposition, but also for informing the broader function of AAA+ proteins in essential cellular processes such as protein degradation, DNA replication, and cytoskeletal reorganization^21,22^.

Here, we combined extensive molecular dynamics (MD) and free energy simulations with an artificial intelligence (AI)–driven Graph Neural Network (GNN) attention model to uncover the mechanisms underlying TnsC filament nucleation and elongation along dsDNA. MD simulations have proven valuable for elucidating the biophysical principles of CRISPR-Cas systems^23–28^, informing strategies for gene knock-in through these systems, and guiding their rational engineering^29,30^. These approaches are now beginning to shed light on the molecular basis of site-specific gene knock-out via RNA-guided transposition^31^. We reveal that ATP plays a pivotal role in initiating TnsC assembly by stabilizing key protein–DNA interactions and promoting DNA remodelling critical for filament formation. Furthermore, our attention-based analysis shows that TnsC preferentially elongates in the 5′→3′ direction, driven by asymmetric dynamics between the DNA-bound and incoming monomers. Together, these findings provide a detailed mechanistic view of TnsC filament formation and offer a foundation for the rational design of programmable, site-specific DNA transposition systems. Finally, our GNN-based attention models establish a new framework for extracting mechanistic information from molecular simulations.

## Results and Discussion

### TnsC alters DNA structure and dynamics

A TnsC filament of approximately 23 nm was based on the ATP-bound structure of the *S. hofmanni* TnsC octamer (PDB 7M99^9^). This filament comprises six TnsC hexamers – each consisting of six monomers – for a total of 36 TnsC subunits arranged in helical turns around dsDNA. The resulting configuration adopts a spiral architecture in line with electron micrographs of the TnsC filament^9^. The system was simulated in explicit solvent reaching a ~15 μs sampling (in three replicates of ~5 μs each). For comparison, MD simulations of canonical B-DNA with an equivalent number of base pairs were also performed.

Analysis of DNA structural dynamics reveals marked differences between TnsC-bound DNA and canonical B-DNA. These disparities are reflected in their root-mean-square deviation (RMSD), spanning a range of ~6 to ~16 Å (Supplementary Fig. 1). The TnsC-bound DNA shows an expansion of the minor groove by ~2 Å and a corresponding narrowing of the major groove relative to B-DNA (**Fig. 2a**), changes often associated with protein-DNA interactions^32,33^. Pronounced DNA kinking and altered twist angles in the TnsC-bound DNA (Supplementary Fig. 2) further contribute to an average of 11.5 base pairs per turn, deviating from the 10.5 in B-DNA, and an increased overall length reaching 233 ± 1.2 Å, with 39.5 ± 0.8 Å per helical turn (**Fig. 2b**, Supplementary Fig. 2).

**Fig. 2.**
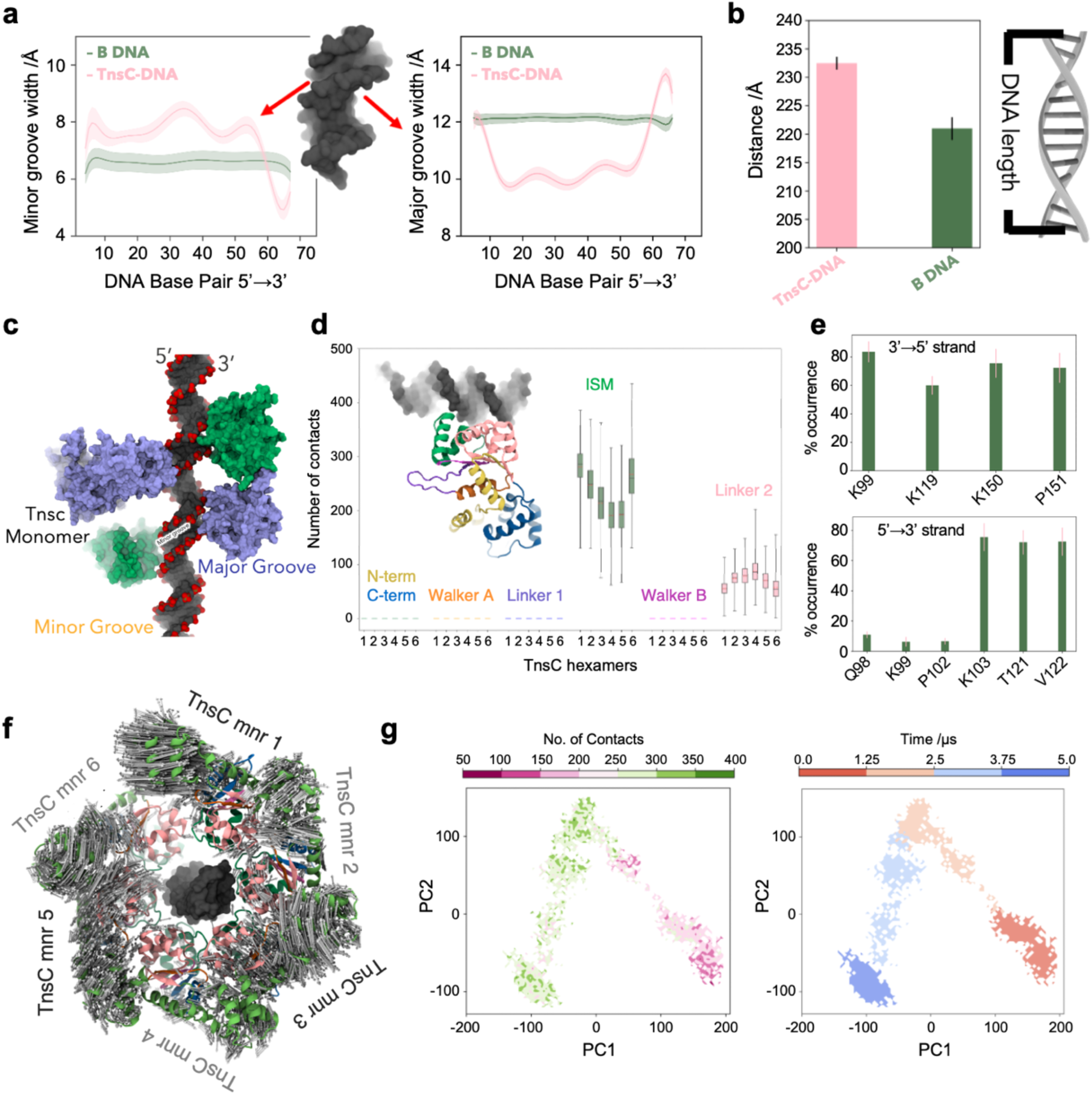
TnsC-induced dynamical remodelling of DNA. **a.** Minor (left panel) and major groove (right panel) widths of the DNA in the TnsC filament (TnsC-DNA, pink) compared to B-DNA (green), from ~15 μs of molecular dynamics simulations. **b.** DNA length, measured as the distance between the centres of mass of the terminal base pairs, shown for TnsC-DNA (pink) and B-DNA (green). Error bars in panels **a** and **b** represent the standard error of the mean from three independent ~5 μs replicates. **c.** Arrangement of TnsC monomers with respect to DNA. For clarity, four representative TnsC monomers (blue and green) are shown to interact with the DNA backbone and the minor groove. **d.** Box plots of the number of contacts between conserved TnsC motifs and the DNA for each hexamer. Contacts were defined as heavy-atom distances ≤ 4 Å between the protein and DNA phosphate backbone, computed over a ~15 μs ensemble. Boxes represent the interquartile range (IQR), with the centre line indicating the median; whiskers extend to the minimum and maximum values within 1.5× IQR. **e.** Statistical distribution (% of the total simulation time) of the contacts between the positively charged/polar residues of the ISM and Walker-B linker regions and the DNA. Error bars are the standard error of the mean from three independent ~5 μs-long simulations. **f.** Porcupine plot of TnsC hexamer dynamics. Top-view of a TnsC hexamer with arrows reflecting the motions along the first principal component (PC). Arrow sizes are proportional to the amplitude of motions. **g.** Conformational sampling of a representative TnsC hexamer along the first and second PCs (i.e., PC1 vs. PC2), plotted with respect to the number of TnsC-DNA contacts (left) and the simulation time (right).

These structural features are consistent with cryo-EM data of the TnsC octamer used as the simulation reference (PDB 7M99^9^), which shows DNA with 11 base pairs per turn and a helical pitch of 40.7 Å (Supplementary Fig. 2c). This agreement indicates that the distortions imposed by TnsC on the DNA within the octamer are preserved along the extended ~23 nm filament. Overall, the DNA undergoes structural adaptations to align with the helical symmetry imposed by the TnsC filament. The positioning of TnsC monomers relative to dsDNA reveals that interactions are primarily localized to the phosphate backbone and the minor groove (**Fig. 2c**), consistent with the observed minor groove expansion (**Fig. 2a**).

Minimum distance analysis (Supplementary Fig. 3a) shows that TnsC monomers are positioned ~3 Å from the phosphate backbone, ~7.5 Å from the minor groove, and ~8 Å from the major groove. An analysis of the DNA contacts with conserved motifs of TnsC was performed for each monomer across the hexameric assemblies (details in Supplementary Methods). We found that the ISM and Linker 2 regions (between the Walker B and C-terminal) establish continuous interactions along the DNA’s length (**Fig. 2d**), underscoring their critical role in maintaining filament stability and DNA binding. Indeed, positively charged residues within these regions persistently interact with the DNA backbone. K99, K119, K150, and P151 maintain steady interactions with the 3′→5′ DNA strand for ~80% of the simulation time, while K103, T121, and V122 primarily engage with the 5′→3′ strand (**Fig. 2e**). Mutational studies have shown that alanine substitutions at K99, K103, and T121 abolish transposition^9,10^, validating the functional significance of our findings.

Principal Component Analysis (PCA) of C⍺ atoms from each hexamer (details in Supplementary Methods) highlights key aspects of TnsC dynamics. A porcupine plot of the first principal component – PC1, i.e., the so-called “essential dynamics” – was mapped onto the 3D structure of a representative hexamer, illustrating the direction and relative amplitude of the collective motions (**Fig. 2f**). The N- and C-terminal regions of each monomer display large-amplitude motions, whereas ISM and Walker B remain relatively stable, reflecting their stronger interactions with DNA. Projections along PC1 and PC2 were plotted against the number of protein–DNA contacts per hexamer (**Fig. 2g** and Supplementary Fig. 4a) revealing that all hexamers traverse the conformational landscape and converge toward states with increased DNA contacts relative to their initial configurations. Together, these findings suggest that the flexible N- and C-termini modulate the dynamics of the ISM region, promoting its effective engagement with DNA.

### ATP initiates TnsC nucleation on dsDNA

Biochemical and structural studies show that TnsC, predominantly monomeric in solution, oligomerizes into helical filaments on dsDNA in an ATP-dependent manner or near the Cas12K– TniQ complex^9^. These filaments recruit the TnsB transposase, yet the molecular mechanisms of TnsC nucleation and its elongation on dsDNA are not understood.

To investigate the nucleation process of TnsC on dsDNA and the role of ATP, we performed free energy simulations. We examined the association dynamics of a representative TnsC monomer with dsDNA in the presence and absence of ATP using the Umbrella Sampling (US) method. Here, a TnsC monomer was initially positioned at a non-interacting distance (~20 Å) from dsDNA, representing the unbound state, while the dsDNA-bound monomer observed in the cryo-EM structure (PDB 7M99)^9^ served as the final state. The process was examined along the RMSD of the protein Cα and the phosphate backbone of the dsDNA nucleotides that interact with TnsC in the cryo-EM structure, relative to the unbound and DNA-bound states (used as the reaction coordinate, RC). The association dynamics of the TnsC monomer with dsDNA was sampled over ~2 μs of converged US runs in the presence and absence of ATP (see the Supplementary Methods, and Supplementary Fig. 5).

After removing biases incurred by the restraints, we computed the free energy landscape as the Potential of Mean Force (PMF). In the presence of ATP, the association process displays a smooth energy profile, reaching a well-defined minimum, corresponding to the DNA-bound state observed in the cryo-EM structure (RC ~2.5 Å; Fig 3a, top graph). By contrast, the apo monomer encounters a higher free energy barrier and fails to reach the cryo-EM-bound configuration, underscoring the critical structural role of ATP in initiating TnsC nucleation on dsDNA. To uncover the structural factors influencing the process, we analysed the ATP-bound and apo monomers at key steps along the RC. Specifically, the initial interaction of TnsC with dsDNA (RC ~4.5 Å, hereafter referred to as “nucleation point”) and the minimum corresponding to the cryo-EM-bound state (RC ~2.5 Å). At the nucleation point, the ATP-bound monomer engages the dsDNA minor groove via the ISM region, followed by a conformational rearrangement that stabilizes the complex at the reference state (RC ~2.5 Å, Fig 3a, top panel). In contrast, in the apo state, the ISM region approaches the dsDNA but fails to reach the reference state (Fig 3a, bottom panel).

**Fig. 3.**
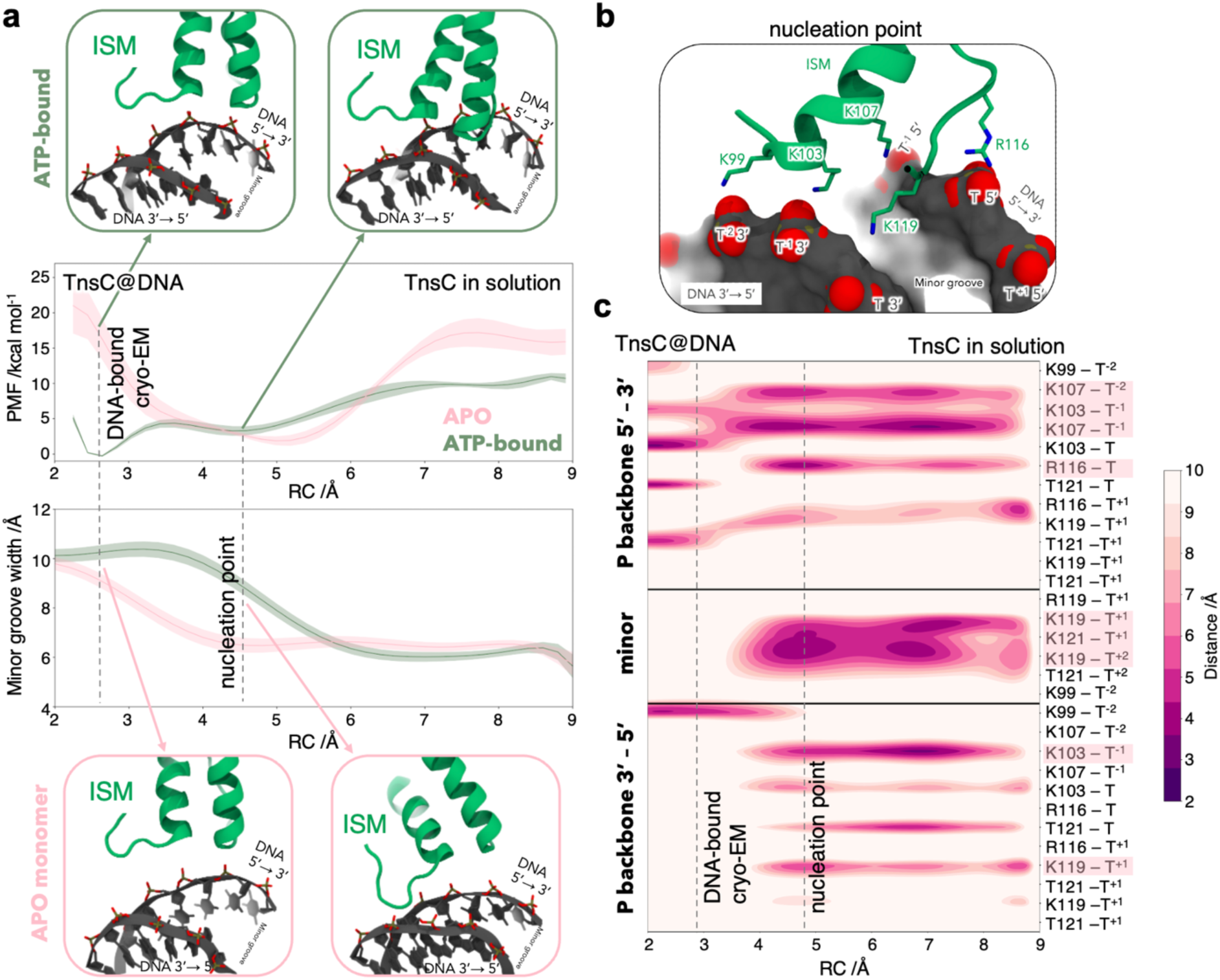
TnsC Nucleation on double-stranded DNA (dsDNA). **a.** Free energy simulations describe the association of a representative TnsC monomer with dsDNA in the presence (green) and absence (pink) of ATP. The free energy profiles (top graph) are computed as the Potential of Mean Force (PMF, kcal/mol) and plotted along the reaction coordinate (RC). The RC is defined as the root-mean-square deviation (RMSD) of the TnsC Cα atoms and dsDNA phosphate backbone atoms interacting with TnsC in the cryo-EM structure (PDB 7M99), relative to the unbound and bound states. The minor groove width (bottom graph) for both the ATP-bound and apo states of TnsC is plotted along the RC. Errors in the free energy profiles are computed via bootstrap error analysis (see Supplementary Methods). Representative snapshots, marked by arrows, correspond to RC values of ~2.5 Å (i.e., the cryo-EM state, with TnsC bound to dsDNA) and ~4.5 Å (i.e., the “nucleation point” of initial TsdC-dsDNA interaction). **b**. b. Close-up view of the interactions formed by the ISM region of TnsC with dsDNA at the nucleation point. **c.** Detailed interactions between positively charged residues within the ISM region and the dsDNA backbone and minor groove along the TnsC–dsDNA association process (i.e., RC). Interactions were quantified as the minimum distances between heavy atoms of ISM side chains and those of the DNA base pairs (see Supplementary Methods).

Notably, in the presence of ATP, the minor groove begins to expand at approximatively the nucleation point and reaches ~10 Å width at the energetic minimum (Fig 3a, bottom graph). In contrast, the minor groove width remains ~7 Å until RC ~3 Å, after which it expands, albeit with an associated increase in energy expenditure. These observations suggest that the ATP-bound TnsC triggers DNA remodelling at the onset of filament formation (i.e., near the nucleation point). It is also noteworthy that throughout the association process, the ATP-bound TnsC displays markedly greater structural stability than the apo monomer (Supplementary Fig. 6b). This suggests that ATP binding stabilizes TnsC, imparting the rigidity needed to engage effectively with dsDNA and promote its remodelling.

We next tracked the interactions between positively charged residues within the ISM region and the dsDNA phosphate backbone along the RC (**Fig. 3b**). In the ATP-bound TnsC, K119 – located within a flexible loop of ISM – engages the minor groove early in the association process (RC ~8.5 Å) and remains anchored through the nucleation point. As the association progresses, K99, K103, K107, and R116 form coordinated contacts with specific phosphate groups, cumulatively stabilizing the complex and culminating in the cryo-EM-like configuration. In contrast, the apo monomer engages dsDNA only at a later stage, as the complex approaches the reference state (Supplementary Fig. 7). These findings indicate that the ISM region initiates filament formation through key positively charged residues, which play a central role in promoting DNA remodelling.

### Mechanism of TnsC filament elongation

Biochemical studies have shown that TnsC filaments assemble around dsDNA exhibiting bidirectional polarity and enabling filament growth in both the 3′→5′ and 5′→3′ directions ^14–16,18^. To probe the mechanisms underlying filament growth and determine whether a directional preference exists, we performed free energy simulations and combined them with a Graph Attention Network (GAT)^34^. Using US simulations, we explored the binding of a second ATP-bound TnsC monomer to a DNA-bound monomer in both the 3′→5′ and 5′→3′ directions (**Fig. 4a**). In each case, the incoming monomer was initially positioned ~20 Å from the DNA-bound monomer, representing the unbound state, with the bound state corresponding to the cryo-EM structure (PDB 7M99)^9^. The RC was defined similarly to that used for the nucleation process (details in Supplementary Methods). US simulations described the binding process through ~2.3 µs of total sampling. The free energy profiles (**Fig. 4a, bottom**) reveal a higher energy barrier for TnsC binding in the 3′→5′ direction compared to the 5′→3′ direction. This elevated transition state suggests that filament elongation is energetically less favorable in the 3′→5′ orientation, indicating a directional preference for 5′→3′ assembly.

**Fig. 4.**
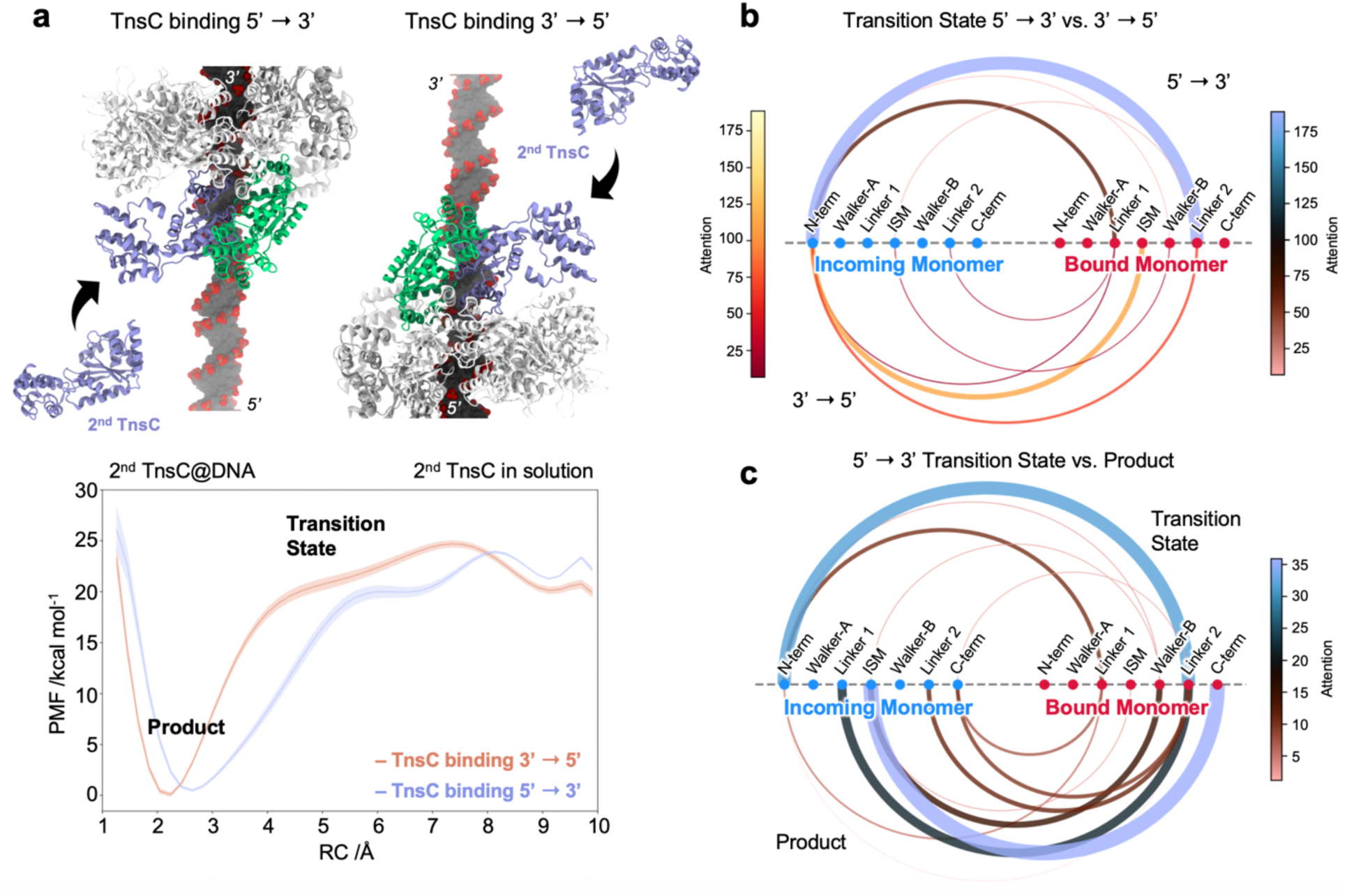
Mechanism of TnsC elongation along double-stranded DNA (dsDNA). **a.** Binding of a second TnsC monomer to a dsDNA-bound monomer in the 5′→3′ (left) and 3′→5′ directions (right). The corresponding free energy profiles (bottom graph), calculated as the Potential of Mean Force (PMF, kcal/mol), are plotted along the reaction coordinate (RC) as defined in Fig. 3 and Supplementary Methods. **b-c.** Graph Attention Network (GAT) models classify conformational states and identify key molecular determinants via attention scores (ATTs). **b.** Arc graphs visualizing the ATTs between the motifs of the incoming and bound TnsC monomers for the transition states in the 5′→3′ (top) and 3′→5′ (bottom) directions, derived from a two-state GAT model. **c.** Arc graphs of the inter-monomer ATTs for the transition (top) and bound (bottom) states in the 3′→5’ direction, obtained from a three-state GAT model. Motif-to-motif interactions are depicted, with arc colour indicating ATT magnitude (see colour scale) and arc thickness representing variance (see main text for details).

To identify the molecular determinants driving filament elongation along the preferred pathway, we developed a Graph Attention Network (GAT) model^34^. GATs are an advanced class of Graph Neural Networks (GNNs)^35,36^ that leverage attention mechanisms to assign adaptive weights to node features, thereby enhancing the identification of critical interaction patterns in graph-structured biomolecular data. Unlike conventional graph-based models that treat all nodes uniformly^37,38^, Graph Attention Networks (GATs) dynamically assign importance to nodes via learned attention scores (ATTs). This mechanism enables GATs to outperform standard GNNs, which uniformly aggregate neighbour information. In our approach, GATs not only classify conformational states but also yield interpretable ATTs that inform the molecular determinants driving classification decisions. Here, each conformational state was encoded as a graph, with nodes (e.g., Cα atoms or residues) annotated with atomic mobility and interaction features. Edges, defined within a 10 Å cutoff, were weighted by inverse inter-residue distances, capturing structural connectivity (details in Supplementary Methods).

The free energy profiles revealed a higher transition state energy barrier in the 3′→5′ direction compared to the 5′→3′ pathway (**Fig. 4a**). To elucidate the biophysical basis of this difference, we developed a two-class GAT model to classify the transition states corresponding to each pathway (**Fig. 4b**). Using this model, we computed attention scores (ATTs) both within and between the incoming and DNA-bound monomers. ATTs were visualized as an arc graph, where the colour scale represents the magnitude of attention and arc thickness corresponds to its variance. High attention scores indicate stable communication or interactions between molecular regions, while high variance highlights dynamic connections that form and break.

This analysis revealed distinct interaction patterns between the transition states of the 5′→3′ and 3′→5′ pathways (**Fig. 4c**), offering mechanistic insights into the differential energetics observed in the free energy profiles. In both pathways, the N-terminal of the incoming monomer primarily communicates with the linker regions of the bound monomer. Notably, the 3′→5′ direction shows an increased number of attentions between the incoming and bound monomers. Analysis of ATT variance reveals greater variability in the interacting regions in the 5′→3′ direction, reflecting enhanced flexibility and dynamic communication at the transition state. Conversely, the 3′→5′ direction exhibits lower ATT variance, indicative of more extensive and stable interactions that restrict conformational dynamics. This suggests that in the 3′→5′ direction, the incoming monomer forms stronger interactions with the bound monomer at the transition state, imposing structural constraints that restrict conformational flexibility. These enhanced inter-monomer contacts provide a mechanistic explanation for the broader transition state window observed in this pathway compared to the 5′→3′ direction (**Fig. 4a**).

To further elucidate the role of interaction dynamics in filament elongation, we separately analyzed intra-monomer ATTs and their variances for the incoming and bound monomers at the transition state in both directions. Notably, in the 5′→3′ direction, the incoming monomer exhibits increased dynamics, as indicated by ATTs with high variance (Supplementary Fig. 8). In contrast, in the 3′→5′ direction, heightened fluctuations are observed in the bound monomer. This suggests a compensatory mechanism at play in TnsC filament elongation. As the filament elongates in the 3′→5′ direction, increased inter-monomer interactions (**Fig. 4b**) restrict the mobility of the incoming monomer (Supplementary Fig. 8) at the transition state, while the bound monomer is more dynamic. This hampers the system to escape the transition state, rendering the 3′→5′ direction less favorable. On the other hand, elongation in the 5′→3′ direction allows greater flexibility in the incoming monomer, facilitating a smoother transition and supporting an energetically more favorable process. Overall, the redistribution of dynamics between monomers – where increased rigidity in one monomer shifts the dynamics to the other – impacts the energetics of filament formation. This dynamic compensation provides a mechanistic basis for the observed preference in filament growth in the 5′→3′ direction.

To gain deeper insight into the mechanism of TnsC filament elongation along the 5′→3′ direction, we developed a three-class GAT classification model that effectively distinguishes unbound, transition state, and bound configurations. Through this model, we computed the ATTs between the incoming and bound monomers at both the transition and DNA-bound states (**Fig. 4c**). Our analysis revealed that, as the incoming monomer approaches the transition state, its N-terminal establishes communication with the bound monomer’s Linker 1 (connecting Walker-A and ISM) and Linker 2 (connecting Walker-B and the C-terminus) regions. Upon reaching the final bound state, a distinct shift in attention patterns emerges. The ATTs primarily link the ISM and Linker 1 regions of the incoming monomer with the Linker 2 and C-terminal regions of the bound monomer. This reorganization reflects a reconfiguration of the interaction network, marking the completion of the binding process and the stabilization of the bound state.

We next computed the intra-monomer ATTs for both monomers in the unbound, transition, and DNA-bound states of the incoming monomer. Notably, the two linker regions and the ISM of the incoming monomer consistently exhibited high ATTs throughout all states, underscoring their structural relevance (Supplementary Fig. 9). Interestingly, the C-terminal progressively decreases its interactions – particularly with the N-terminal α-helix and ISM – as the incoming monomer transitions from the unbound to the transition and final bound states. This indicates that the C-terminal becomes structurally disengaged, suggesting its increased availability for interacting with the bound monomer. This trend is indeed captured in the ATTs derived from the three-state GAT model characterizing interactions between the bound and incoming monomers (**Fig. 4c**). Conversely, the bound monomer displays relatively moderate changes in ATT scores across the unbound, transition, and product states, reflecting a conserved structural framework throughout the process. Notably, the C-terminal region consistently exhibits low ATT values in all states, indicating it undergoes minimal conformational rearrangement compared to the incoming monomer. Overall, these attention patterns indicate that the incoming monomer undergoes flexible restructuring, especially in its C-terminal region, enabling it to interact with the relatively rigid and pre-formed structure of the bound monomer. This differential behavior may be essential for driving directional binding and complex formation

## Conclusions

In summary, this study provides a mechanistic framework for understanding the nucleation and elongation of TnsC filaments in type V-K CRISPR-associated transposons (CAST), revealing how ATP binding and DNA remodeling synergize to initiate and propagate filament formation along dsDNA. Using molecular and free energy simulations, we demonstrate that ATP plays a pivotal role in stabilizing key interactions – particularly through the initiator-specific motif (ISM) – that are essential for DNA remodeling and initiate the nucleation of the TnsC monomers on dsDNA. Moreover, by introducing a Graph Neural Network (GNN)-based attention model, we uncover a dynamic, directionally biased mechanism of filament elongation in the 5′→3′ direction. We demonstrate that the structural flexibility is asymmetrically distributed between incoming and DNA-bound TnsC monomers to facilitate directional assembly, favoring the 5′→3′ direction. This interplay of dynamics, energetics, and structural transitions not only explains the preferential growth of TnsC filaments but also introduces a model of dynamic compensation during monomer recruitment. Furthermore, our GAT models enable the extraction of mechanistic insights from molecular simulations by assigning dynamic, context-specific attention weights to molecular interactions. These models incorporate predictive relevance directly into the analysis of structural and dynamical features. This approach not only enhances interpretability but is also readily transferable across diverse biological systems. Together, these findings bridge atomic-level structure with function in CAST systems and open new avenues for engineering programmable DNA integration platforms with improved specificity and control.

## Materials and Methods

### Molecular Dynamics simulations

Molecular simulations were conducted on an ATP-bound TnsC filament model based on the 3.20 Å cryo-EM structure (PDB: 7M99)^9^, comprising eight TnsC monomers wrapped around dsDNA. A ~240 Å TnsC filament was generated, including six hexameric rings encircling 70 bp of dsDNA. A canonical B-form DNA model of equivalent length was also generated. Both systems were solvated in periodic water boxes and neutralized with counterions. Molecular dynamics (MD) simulations were performed using a protocol tailored for protein/nucleic acid complexes^39^, also applied in studies of CRISPR-Cas systems^24,26,29,31^. The Amber ff19SB^40^ force field was employed, incorporating the OL15^41^ corrections for DNA. The TIP3P^42^ model was used for water molecules. An integration time step of 2 fs was applied. For each system, ~15 µs of total ensemble sampling was obtained from three independent MD runs of ~5 µs each, initiated from different starting coordinates and velocities. Full details on the simulation protocol are reported in the Supplementary Methods.

### Umbrella Sampling Simulations

Umbrella sampling (US)^43^ simulations were employed to compute the free energy profiles for two key steps in TnsC filament polymerization: (i) nucleation, defined as the binding of a representative TnsC monomer to DNA with and without ATP; and (ii) elongation, modeled as the binding of a second TnsC monomer to the monomer– DNA complex, in both the 5′→3′ and 3′→5′ directions. Initial configurations for the US simulations were prepared by placing TnsC monomers ~20 Å from the dsDNA to represent a non-interacting, unbound state, while the cryo-EM structure of the dsDNA-bound TnsC (PDB: 7M99) was used as the bound state reference. To describe the nucleation process, the reaction coordinate (RC) was defined as the root-mean-square deviation (RMSD) of the Cα atoms of a TnsC monomer and the phosphate backbone of the interacting dsDNA segment (PDB 7M99: Chain I, nucleotides 6–14; Chain J, nucleotides 8–16), relative to the unbound and cryo-EM-bound states. For the elongation process, the RC was similarly defined as the RMSD of the Cα atoms of both TnsC monomers and the phosphate backbone of the interacting dsDNA region (Chain I, nucleotides 3–14; Chain J, nucleotides 8–19), relative to both states. These RCs enabled to account for the association of the TnsC monomers to dsDNA and the conformational changes of the protein. US simulations for the nucleation process were conducted across 46 overlapping windows, with the RC ranging from ~10 Å to 0.2 Å. For elongation, 56 windows were used, spanning ~11 Å to 0.2 Å. A harmonic restraint with a spring constant of 50 kcal/mol·Å^2^ was applied, and window spacing was set to 0.2 Å to ensure adequate overlap of probability distributions. For each system, ~20 ns trajectories were obtained per US window, resulting in ~1 µs of collective sampling per system (and a total of ~4 µs). Statistical uncertainties in the free energy profiles were estimated using Monte Carlo bootstrap analysis (full details in the Supplementary Methods).

### Graph Attention Network

Graph Attention Networks (GAT)^34^ represent an advanced variant of Graph Neural Networks (GNNs) ^35,36^ that utilize attention mechanisms to selectively weigh node features, enhancing the learning process from graph-structured data. Here, our molecular systems are represented as graphs, where nodes (e.g., atoms or residues) are characterized by various features such as mobility and interaction properties, while edges define their connectivity, capturing structural and functional relationships.

A residue-level graph *G*(*V*, *E*) was constructed for each frame in the MD trajectory, where each residue’s Cα atom was represented as a node *ν_i_* ∈ *V*, and edges *e_ij_* ∈ E were defined between residue pairs located within a 10 Å cutoff. Edge weights *w_ij_* were calculated as the inverse of the Euclidean distance between the corresponding Cα atoms, i.e., 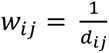 for *d_ij_* ≤ 10Å. Each node was annotated with features *x_i_*, including structural and dynamic descriptors, and a GAT was trained to classify each frame into one of three conformational states: reactant, transition state (TS), or product.

The network architecture consisted of stacked *GATConv* layers ℓ, each computing attention weighted node embeddings followed by *ReLU* activation. The final layer output was aggregated using a global mean pooling operation across all nodes in the graph, and the resulting graph-level embedding was passed through a linear classification layer followed by a *Softmax* activation. This process can be expressed as:

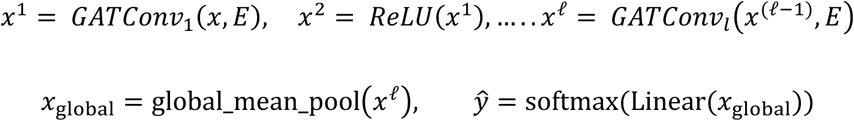

Model training was performed using the cross-entropy loss function:

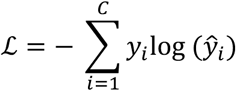

where *C* = 3 is the number of classes, *y_i_* is the true label, and *ŷ_i_* is the predicted probability. The dataset was divided into training (90%) and test (10%) sets using stratified sampling to preserve class proportions. Early stopping was employed based on validation loss to mitigate overfitting.

GAT employs an attention mechanism to learn the relative importance of residue-residue interactions. For each node pair (*i*, *j*), an attention coefficient *α_ij_* is computed as:

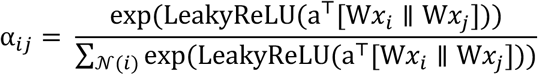

where W is a learnable transformation matrix, a is the attention vector, || denotes concatenation, and 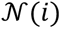 represents the neighborhood of node *i*. This formulation enables the model to integrate structural (topology), dynamic (node features), and contextual (local environment) information.

The final output depends on the attention-weighted aggregation of node features through all layers, such that the model’s predictions are directly influenced by the cumulative contributions of residue-level interactions. Attention scores from multiple layers were summed to obtain a combined score:

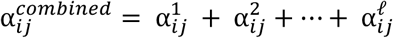

These combined attention scores were used to assess inter-residue communication, particularly focusing on interactions between residues from distinct monomers. Average attention per residue pair was computed across all frames:

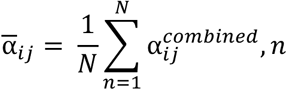

where *N* is the number of frames. Variability in these attention scores was calculated as:

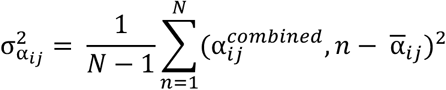

A high value of 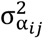 indicates that the importance of a particular residue-residue interaction changes substantially across different structural states, suggesting the interaction is transient or state-dependent. Conversely, consistently high or low average attention values across states highlight stable interactions that are persistently important or consistently negligible, respectively. These attention-based metrics capture facets of residue communication that elude traditional structural fluctuation analyses, such as RMSF or covariance matrices, which do not directly integrate with predictive modelling tasks.

To further enhance interpretability, residues were grouped into functionally annotated regions (e.g., Walker motifs, insertion sequences), and attention was aggregated at the region level. The total attention between regions *R*1 and *R*2 was computed as:

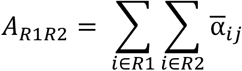

with frame-wise variance computed analogously:

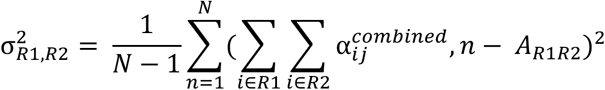

These regional metrics offer robust and interpretable summaries of inter-domain communication patterns and their modulation across conformational transitions.

In contrast to conventional MD analysis techniques, which rely primarily on structural deviations or pairwise correlations, the attention-based framework incorporates predictive relevance directly into the interaction analysis. By evaluating both average attention and its variance, this method identifies residue interactions/connections that are stable, transient, or state-specific, thereby providing a more comprehensive view of conformational dynamics relevant to function. This approach offers an integrative framework to decode mechanistically informative features from MD simulations using deep learning-based graph representations.

### Analysis of structural data

Evaluation of the DNA structural dynamics involved the assessment of key structural parameters along the MD simulations. The widths of the minor and major grooves were analysed according to the definition by Suzuki & Yagi^44^, by measuring the shortest inter-strand phosphate distances. DNA bending and twisting were measured, according to previous computational studies of DNA dynamics^45,46^ (details in the Supplementary Methods). The length of the DNA was determined as the distance between the centers of mass of its first and last base pairs. The interactions between TnsC and the DNA were evaluated by calculating the minimum distances between the heavy atoms of the protein and the DNA phosphate backbone, measured separately for the 5′→3′ strand and the 3′→5′ strand, as well as the minor and major grooves heavy atoms. Contact analysis was performed to compute the number of contacts between the DNA and the conserved motifs of TnsC (i.e., Walker-A, Walker-B, ISM, Linkers 1-2, and the N- and C-termini). Contacts were considered formed when the distance between the heavy atoms of the protein and the DNA phosphate backbone is ≤ 4 Å. This analysis was performed for each TnsC monomer across the hexameric assemblies in the TnsC filament. The analyses were performed on a collective ensemble of ~15 μs per system, obtained from three independent ~5 μs replicates. Error estimates were calculated as the standard error of the mean across the three replicates. Full details are reported in the Supplementary Methods.

## Acknowledgments

This material is based upon work supported by the NIH (Grant No. R01GM141329 to G.P.) and the NSF (Grant No. CHE-2144823 to G.P.). GP acknowledges support by the Alfred P. Sloan Foundation (Grant No. FG-2023-20431) and the Camille and Henry Dreyfus Foundation (Grant No. TC-24-063). The computational studies performed here were carried out using Expanse at the San Diego Supercomputing Center through allocation MCB160059 and Bridges2 at the Pittsburgh Supercomputer Center through allocation BIO230007 from the Advanced Cyberinfrastructure Coordination Ecosystem: Services & Support (ACCESS) program, which is supported by NSF support grants #2138259, #2138286, #2138307, #2137603, and #2138296.

## Author Contribution

CP and MA performed molecular simulations, analysed the data, and contributed to the development of the Graph Attention Network method for analysing molecular dynamics simulations. MA and SS implemented the Graph Attention Network code. CP drafted the initial manuscript. GP conceived the project, supervised the research, and wrote the final manuscript with critical input from all authors.

## Data Availability Statement

The Graph Attention Network code is available from the corresponding author upon request.

## Additional Information

Additional information is available as a supplementary information.

